# Assessing the multimodal tradeoff

**DOI:** 10.1101/2021.12.08.471788

**Authors:** A. Sina Booeshaghi, Fan Gao, Lior Pachter

## Abstract

Single-cell and single-nucleus genomics assays are becoming increasingly complex, with multiple measurements of distinct modalities performed concurrently resulting in “multimodal” readouts. While multimodal single-cell and single-nucleus genomics offers the potential to better understand how distinct cellular processes are coordinated, there can be technical and cost tradeoffs associated with increasing the number of measurement modes. To assess some of the tradeoffs inherent in multimodal assays, we have developed snATAK for preprocessing sequencing-based high-throughput assays that measure single-nucleus chromatin accessibility. Coupled with kallisto bustools for single-nucleus RNA-seq preprocessing, the snATAK workflow can be used for uniform preprocessing of 10x Genomics’ Multiome and single-nucleus ATAC-seq, SHARE-seq, ISSAAC-seq, spatial ATAC-seq and other chromatin-related assays. Using snATAK, we are able to perform cross-platform comparisons and quantify some of the tradeoffs between Multiome and unregistered single-nucleus RNA-seq/ATAC-seq experiments. We also show that snATAK can be used to assess allele concordance between paired RNAseq and ATACseq. snATAK is available at https://github.com/pachterlab/snATAK/.

## Introduction

Single-cell and single-nucleus genomics assays have greatly expanded in scope in recent years, especially in terms of the ability to make multiple simultaneous measurements in single-cells (“Method of the Year 2019: Single-Cell Multimodal Omics” 2020). Multimodal genomics assays are providing increasingly resolved views of the molecular biology of the cell, but improvements in breadth of measurement come with tradeoffs in data quality and cost. For example, the ECCITE-seq assay (Mimitou et al. 2019), which offers the potential for detecting surface proteins alongside single-cell RNA measurements, for assessing clonotypes, provides information on multiple modalities but at the cost of less genes detected per molecule sequenced than with single modality single-cell RNA-seq ((Gorin, Svensson, and Pachter 2020). One group of assays for which the multimodal tradeoff is particularly important to assess are those providing simultaneous measurement of transposase-accessible chromatin (ATAC-seq) and gene expression (RNA-seq) in single nuclei (Yan et al. 2020). Following the introduction of single-nucleus ATAC-seq in (Buenrostro et al. 2013) and multimodal single-nucleus RNA-seq and ATAC-seq (Cusanovich et al. 2015), there are now numerous variants and multimodal extensions (Ma et al. 2020; Xu et al. 2022) that have proved to be valuable for biological discovery (Ogbeide et al. 2022; Buenrostro et al. 2015). Despite the importance of these assays, there has been little assessment of the performance and cost tradeoffs incurred when using them.

The assessment of multimodal tradeoffs across assays and technologies requires uniform data processing, and while there have been some attempts to standardize processing for single-cell RNA-seq (Melsted et al. 2021; Battenberg et al. 2022) and single-nucleus ATAC-seq (Yu et al. 2020; Chen et al. 2019), many multimodal assays are currently published with standalone custom workflows (Wu et al. 2022; Ma et al. 2020; Cheow et al. 2016; Healey, Bassham, and Cresko 2022). In particular, the raw data produced in the ATAC-seq portion of multimodal experiments consists of large numbers of reads, whose preprocessing to identify “peak” regions and counts can pose formidable challenges. One problem is that different assays lead to distinct and complex read structures, making it difficult to perform cross-platform comparisons or to combine datasets (Sina Booeshaghi, Chen, and Pachter 2023). An additional problem is that reads for barcodes obtained by assaying chromatin accessibility must be aligned to the genome while reads for barcodes obtained from assaying transcription must be aligned to the transcriptome all while performing concordant barcode error correction between the modalities. “Turnkey” solutions, such as 10x Genomics’ Cell Ranger ATAC for processing Chromium based snATAC-seq or Cell Ranger ARC for 10x Genomics Multiome preprocessing, while offering specialized preprocessing of multimodal data, can require long runtimes, depend on large memory requirements, and does not generalized across technologies hindering the development of reproducible workflows for data analysis.

In previous work, we presented a modular and efficient approach to single-cell RNA-seq (scRNA-seq) and single-nucleus RNA-seq (snRNA-seq) preprocessing that combines the pseudoalignment program kallisto (Bray et al. 2016) with a suite of tools called bustools (Melsted et al. 2021). These tools facilitate the development of highly efficient and modular workflows for scRNA-seq and snRNA-seq preprocessing that are easy to run using a wrapper called kb (Melsted et al. 2021). Recently, Giansanti et al. (Giansanti, Tang, and Cittaro 2020) developed a pseudoalignment approach for ATAC-seq based on kallisto pseudoalignment of reads to predefined DNAseq hypersensitive sites. With their workflow, they were able to produce results in-line with standard results, but much faster with a substantially smaller memory footprint. Most importantly, because kallisto can process reads from any technology thanks to customizability afforded by the “technology string” input, a kallisto-based ATAC workflow makes possible uniform preprocessing of data from different technologies. Motivated by (Giansanti, Tang, and Cittaro 2020), we have developed snATAK, which incorporates kallisto, bustools, and several other tools in a workflow that facilitates the preprocessing of snATAC-seq data from numerous technologies in minimal computing environments such as Google Colaboratory without the need for pre-defined genome regions (see Methods, Figure1a).

**Figure 1:**
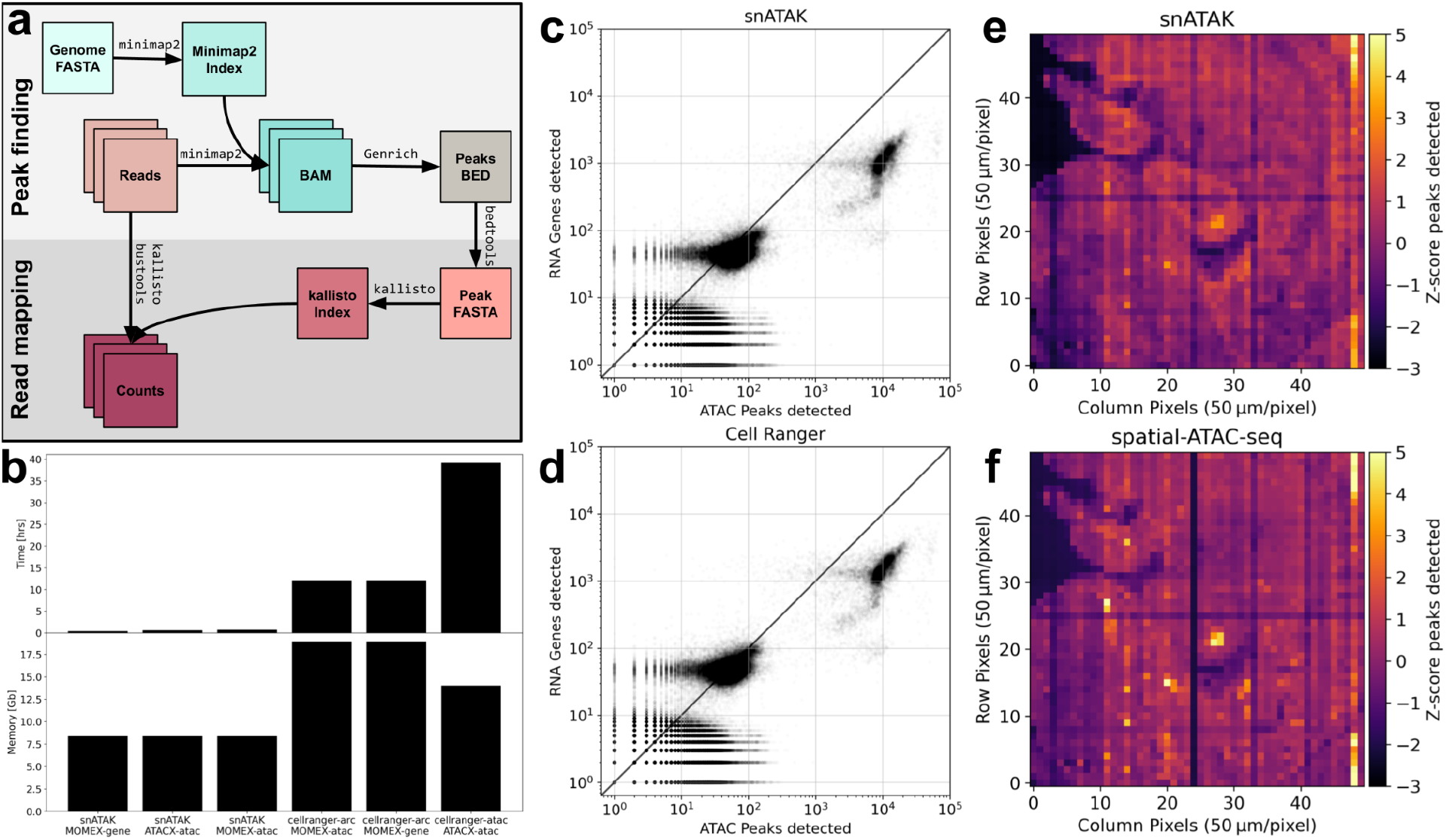
(a) Overview of the snATAK workflow. Reads are first aligned to the genome using Minimap2. Peaks are then called with Genrich and a kallisto pseudoalignment index is constructed from the resultant peaks. Peak counts are obtained using kallisto bustools. (b) Runtime and memory comparisons between snATAK and Cell Ranger ATAC and Cell Ranger ARC on the 10kPBMC datasets assayed with 10x Multiome and 10x snATACseq. (c, d) A comparison of snATAK quantification to Cell Ranger ARC quantification on 10k PBMC cells assayed with the 10x Genomics Multiome kit. For each cell the number of genes detected is plotted against the number of peaks detected. (e, f) A comparison of snATAK quantification to Cell Ranger ATAC v1.2 quantification on a spatial ATAC-seq dataset. Row-column pairs correspond to pixels and are colored by the z-score of the number of peaks detected.

## Results

To validate the suitability of snATAK to assessing multimodal tradeoffs, we first compared results of preprocessing with snATAK to those obtained from two different analysis workflows applied to data from two different technologies: Cell Ranger ARC preprocessing of 10x Genomics Chromium Multiome (Figure 1b,c), and Cell Ranger ATAC v1.2 preprocessing of spatial-ATAC-seq (Figure 1d,e). A comparison of snATAK results to Cell Ranger ARC results on the multimodal detection plot (Figure 1b,c) showed the results to be nearly identical for both the RNA and ATAC data, with the only notable difference being slightly fewer ATAC peaks detected with Cell Ranger ARC than with snATAK. Similarly, a processing of spatial ATAC-seq data from (Deng et al. 2022) with snATAK produced similar results to those obtained by the authors with a workflow based on Cell Ranger ATAC v1.2, although column 42 in the heatmap produced from Cell Ranger ATAC v1.2 was visibly lower than what was obtained with snATAK (see Methods). snATAK produces concordant results with Cell Ranger ARC and Cell Ranger ATAC while running much faster and using less memory (Figure 1b). Whereas snATAK takes 22 minutes to process multiome RNA and 35 minutes to process multiome ATAC, Cell Ranger ARC takes 12 hours for both of these modalities.

Given that snATAK preprocessing produces results concordant with the standard and most widely used tools for single-nucleus ATAC and scRNA-seq preprocessing, we proceeded to explore the relationship between 10x Genomics snRNA-seq, 10x Genomics snATAC-seq, and 10x Multiome that produces both measurements of transcription and chromatin accessibility in nuclei. Notably, the software 10x Genomics provides for analysis of these different data types is not the same; for example, Cell Ranger ARC is used for Multiome whereas Cell Ranger ATAC is used for 10x Genomics single-nucleus ATAC-seq. Despite these differences, a comparison of the ATAC portion of 10x Multiome to ATAC seq across different 10x Genomics machines suggests assay-specific variability that we sought to investigate (Supplementary Figures 1,2).

In a direct comparison of 10x Multiome to 10x snATAC-seq on PBMCs (Figure 2a; see Methods), we found two differences: The Multiome-ATAC contained a large number of multiplets (extra bump in the knee plot) and fewer peaks detected for biologically relevant nuclei (nuclei between the two dashed lines in Figure 2a). The extra multiplets in the Multiome experiment are probably due to overloading; in the experiment ∼16,100 nuclei were loaded resulting in 10,974 nuclei for downstream analysis. The biologically relevant nuclei assayed with snATACseq yielded an average of 9,833 peaks detected while the Multiome-ATAC yielded an average of 8,508 peaks detected. The cells assayed with scRNAseq yielded an average of 5,824 UMIs while the nuclei assayed with the Multiome-RNA yielded an average of 2,481 UMIs. Overall, the number of reads counted per peak was similar between the Multiome and snATAC-seq. In a comparison of the Multiome-RNA to scRNAseq (Figure 2c) we found that there is an efficiency tradeoff, with fewer UMIs identified per read with Multiome than with scRNA-seq, suggesting that Multiome-RNA libraries are less complex than scRNA-seq libraries. Together, these results show that there is a tradeoff between Multiome vs. single-modality single-nucleus assays. On the other hand, 10x Genomics Multiome provides chromatin accessibility and gene expression data that is registered to the same cells, greatly simplifying the task of connecting chromatin changes to gene expression changes in cell types (Lee, Kaestner, and Li 2023).

**Figure 2:**
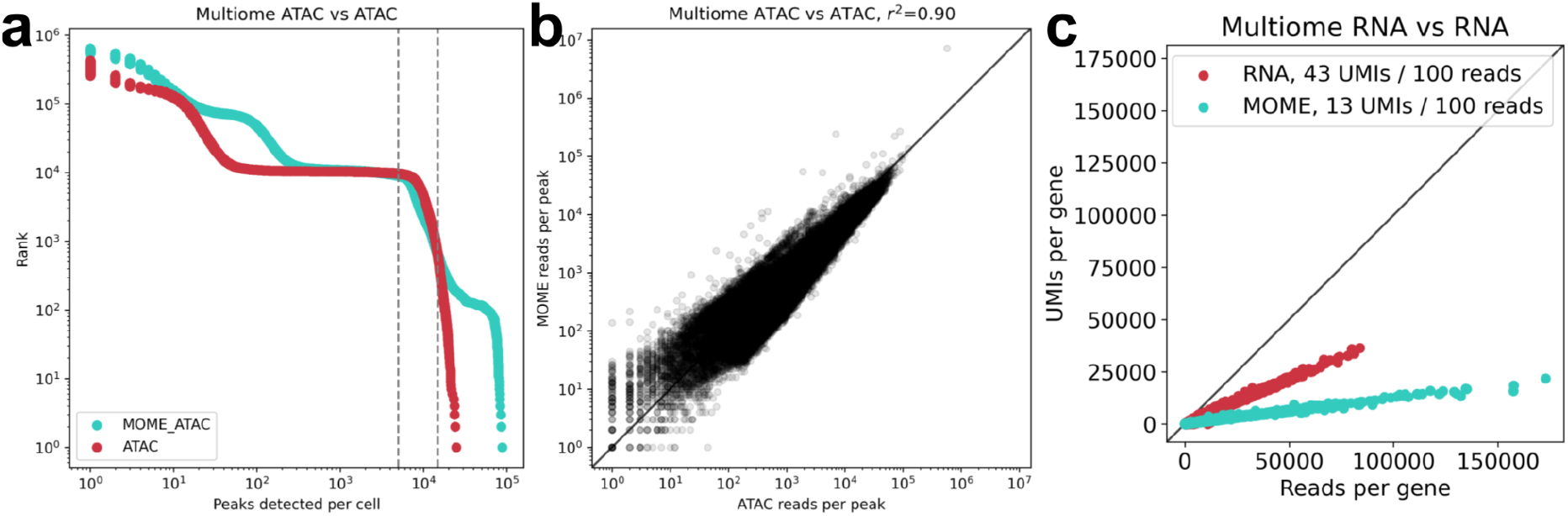
(a) Knee plots of peaks detected in 10x snATACseq (red) and 10x Multiome-ATAC (blue). Dashed lines indicate threshold for putative nuclei. (b) A scatterplot showing the number of reads per peak in 10x Multiome and 10x snATAC-seq data quantified with snATAK. (c) The number of UMIs per 100 reads per barcode for cells assayed with 10x scRNAseq and nuclei assayed with 10x Multiome-RNA.

The uniform processing of data from different technologies that is facilitated by snATAK can also be used to compare technologies to each other to assess whether multimodal tradeoffs are technology specific (Figure 3, Supplementary Figure 3). We compared snATAK processed mixed human HEK293T and mouse 3T3 cell lines assayed with ISSAAC-seq (Xu et al. 2022) to the same mixed cells assayed with SHARE-seqV2 (Ma 2022; Ma et al. 2020). To make a direct comparison, the datasets were subsampled to the same number of both RNA and ATAC reads prior to preprocessing with snATAK (see Methods). The SHARE-seqV2 assay did not perform equally well on human and mouse, with many fewer mouse nuclei yielding high quality data than human (394 vs. 1,159). With the same depth of sequencing, ISSAAC-seq was able to recover 2,998 human cells and 3,556 mouse cells. Even adjusting for multiplets, which were more common in ISSAAC-Seq than in SHARE-seqV2 as the former is a droplet-based rather than well-based method, ISSAAC-seq produced a lot more high quality cells than SHARE-seqV2.

**Figure 3:**
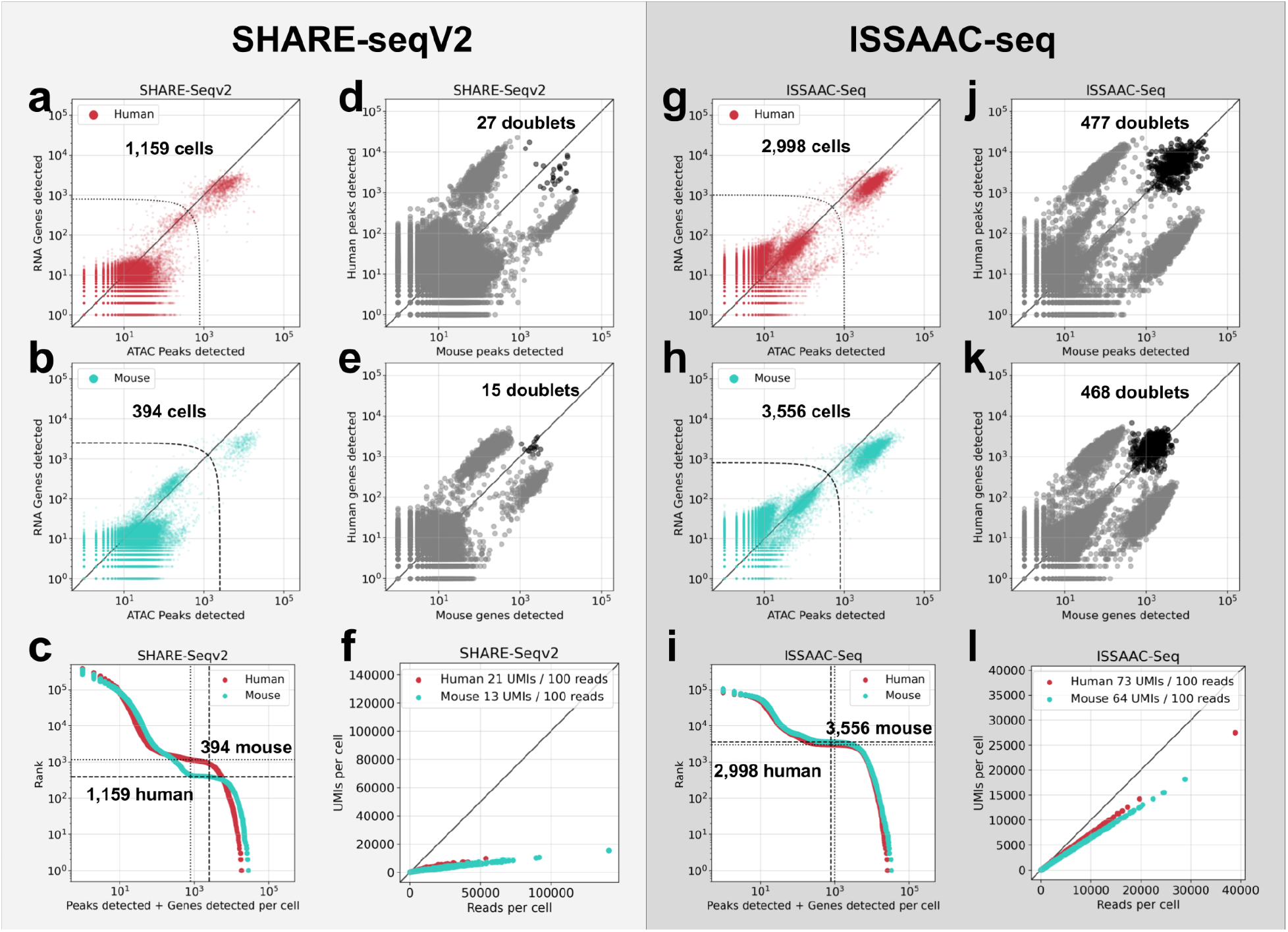
(a,b,c) SHARE-seqV2 RNA genes detected and ATAC peaks detected in human and mouse, and a knee plot computed on the sum of genes and peaks detected. (d,e) SHARE-seqV2 barnyard plots on a log-log axis (clawplots) revealing multiplets. (f) Yield of RNA molecules as a function of reads sequenced. (g,h,i) ISSAAC-seq RNA genes detected and ATAC peaks detected in human and mouse and a knee plot computed on the sum of the genes and peaks detected. (j,k) SHARE-seqV2 barnyard plots on a log-log axis (clawplots) revealing multiplets. (l) Yield of RNA molecules as a function of reads sequenced

The poor SHARE-seqV2 results translate to a much higher cost: ISSAAC-seq yielded 73 human and 64 mouse molecules per 100 reads sequenced, versus 21 human and 13 mouse UMIs per 100 reads sequenced with SHARE-seqV2. However differences between the assays aside, we found that they both exhibit a tradeoff relative to unimodal single-cell RNA-seq. Both SHARE-seqV2 and ISSAAC-seq display much higher technical variation than 10x Genomics single-cell RNA-seq (Supplementary Figure 4). This increase in technical variance, ranging from a factor of 3 to 6, means that with multimodal RNA and ATAC assays it is harder to distinguish biological signal from noise, e.g. in identifying differential genes between conditions. Our analysis demonstrates the utility of uniform processing with snATAK for identifying tradeoffs arising from the choice of technology for multimodal single-nucleus genomics.

One of the benefits of multimodal single-nucleus RNA and ATAC-seq is the ability to directly associate allele-specific expression with strand-specific open chromatin. The snATAK workflow, by virtue of aligning reads to a custom index constructed with kallisto, makes possible genotyping via alignment of reads to pairs of regions around SNPs containing the reference and alternate alleles respectively (see Methods). Using this approach, we genotyped cells in the 10x Multiome PBMC dataset, and computed allele-specific RNA counts together with strand-specific ATAC counts at heterozygous SNPs in nuclei. At SNPs inside genes, where ATAC reads indicated a strong strand preference across cell type, gene expression was strongly allele specific in concordance with the open chromatin strand (see Methods, Figure 4a). While cell type effects were readily measurable, multimodal RNA and ATAC cannot currently be used to resolve allele-specificity in individual cells (Figure 4b). We replicated these results in a Bone Marrow sample assayed with SHARE-seqV2 (Figure 4c,d). Taken together, the results in

**Figure 4:**
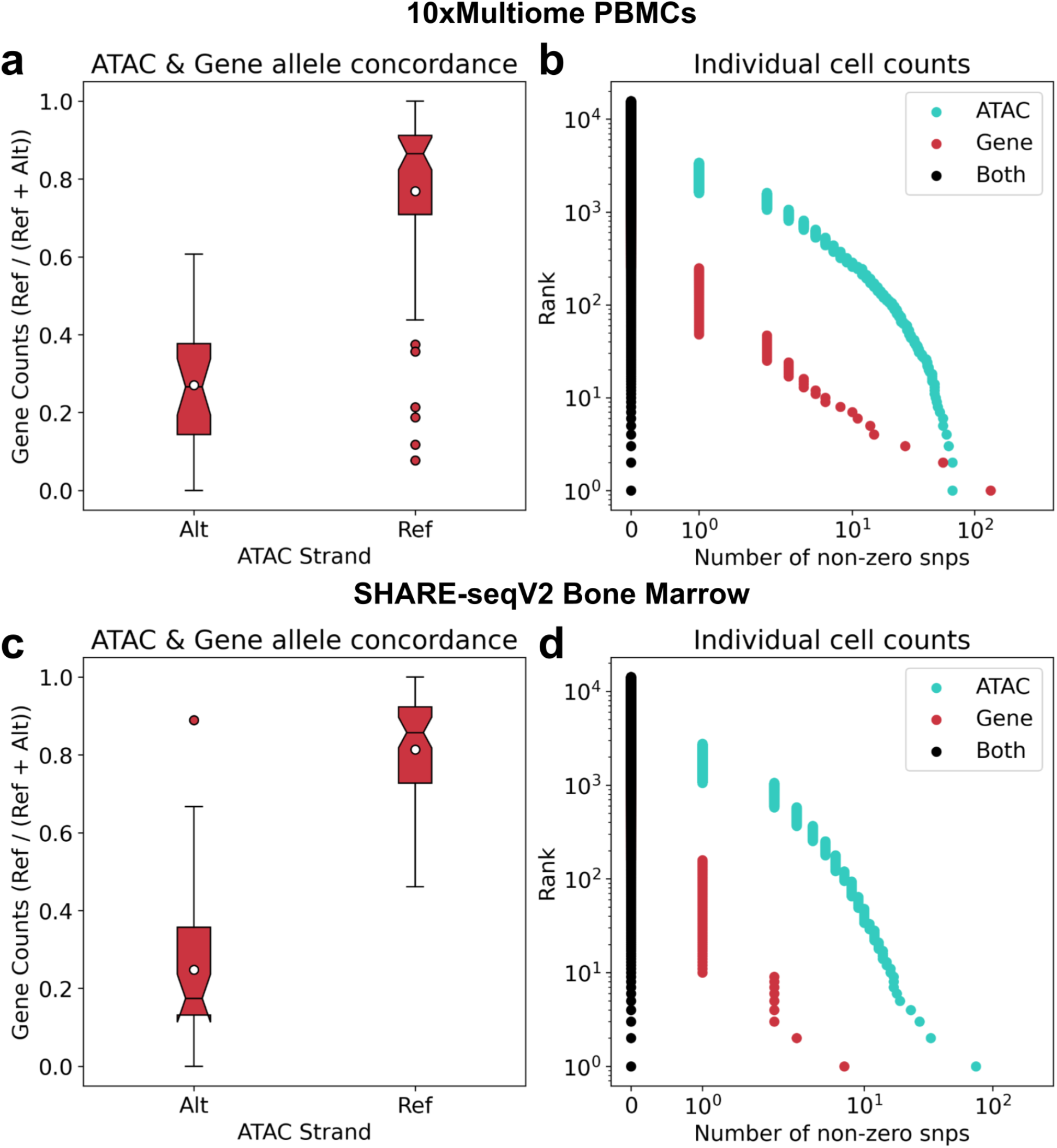
(a) Allele usage concordance between gene expression and open chromatin in PBMCs. Each point is a cell type-SNP combination. (b) Allele-specific counts in single-nuclei in PBMCs assayed with 10x Multiome. (c,d) Same as (a,b) except in human bone marrow assayed with SHARE-seqV2.

Figures 4a,c vs. Figures 4a,d provide an existence proof that RNA and ATAC are allele concordant at the single-cell level, without providing evidence of this in any given individual cell.

## Discussion

Rapid developments in single-cell genomics technologies have been accompanied by an explosion in assay-specific data and processing requirements. Alongside the deluge of data, numerous software tools for data analysis have been developed (Zappia, Phipson, and Oshlack 2018) as well as custom scripts and new programs to process data from almost every new assay. As a result, analysis of datasets published by different groups, possibly with different technologies, is challenging and currently requires extensive “batch correction”, to account for computational variability, a process that can be problematic (Tyler, Bunyavanich, and Schadt 2021). While the snATAK workflow does not solve the problem of how to merge datasets, by virtue of offering a solution for uniform processing of snRNA-seq, snATAC-seq, and Multiome data, it removes technical variability introduced by preprocessing softwares. More generally, the methodology we have presented can facilitate analysis of tradeoffs associated with a wide variety of multimodal assays, and should assist researchers in making informed decisions about experimental design.

While we have demonstrated use of the snATAK workflow for preprocessing snRNA-seq, snATAC-seq, and Multiome assays, the method is suitable for a wide variety of other assays. Essentially any assay that requires quantification of genome alignments can be preprocessed with snATAK. For example, in (Gao and Pachter 2021) we showed that the method can be used for preprocessing Hi-C data as well. Moreover, as interest in snATAC-seq continues to grow (Baek and Lee 2020), there is an increasing need for efficient and accurate preprocessing and analysis software that can facilitate reproducible workflows.

The snATAK workflow is suitable for tasks that are difficult to perform with current methods. For example, we have shown that snATAK can be used for allele-specific analysis of multimodal data, even in the absence of genotype data. Our demonstration of cell-type allele-specificity in both gene expression and open chromatin suggests that single-cell eQTL studies using multimodal experiments are likely to yield interesting results.

Our assessment of the tradeoffs in performing multimodal experiments hints at general tradeoffs that may be worthwhile considering for all multimodal single-cell genomics experiments. In particular, it appears that library complexities vary across technologies, inducing considerable cost differentials. A key question is to what extent the tradeoffs we’ve identified affect downstream analysis results, but a thorough investigation of this is beyond the scope of this paper. However, despite the challenges of multimodal genomics, our work shows that uniform preprocessing is possible, and there is hope for large-scale integrative analysis of the vast amounts of public genomics data that is being generated.

## Acknowledgements

We thank Xun Wang for helpful suggestions. The work was possible thanks to support by the Beckman Institute at Caltech for the Caltech Bioinformatics Resource Center. F.G. and L.P. were supported in part by NIH R01 DK126925-01. A.S.B. and L.P. were supported in part by NIH 5UM1HG012077-02.

## Methods

### snATAK workflow

Our approach consists of first mapping reads to a reference genome using Minimap2 (Li 2018). This tool was selected for its low memory footprint, enabling snATAK to run on Google Colab, thus facilitating usability and reproducibility. Following genome mapping, snATAK identifies putative open chromatin regions with Genrich. A BED file of putative “peaks” is then transformed into a FASTA file of peak sequences using bedtools. A kallisto pseudoalignment index is made from the FASTA file and reads are remapped using kallisto. The purpose of this step is threefold: the pseudoalignment is robust to multi-mapping reads, re-mapping can be variant specific, and the kallisto bustools infrastructure lends itself naturally to counting read pileups. Thus, the Minimap2 step serves as to identify the relevant portions of a reference genome for further preprocessing, and then the snATAC-seq preprocessing is tackled with existing infrastructure borrowed from single-nucleus RNA-seq (Figure 1a). The snATAK output is compatible with the Signac (Stuart et al. 2021) and ArchR (Granja et al. 2021) packages.

### Downloading data SHARE-seqV2 and ISSAAC-seq data

The SHARE-seqV2 human-mouse and bone marrow datasets used to demonstrate the SHARE-seqV2 processing software (Github) and (Hu et al. 2023) were downloaded from the GEO (GSE207308). The single-nuclei RNA-seq reads for the human-mouse and BMMC dataset were downloaded with ffq (Gálvez-Merchán et al. 2022) and wget from GEO accession GSM6284347 and GSM6284350 respectively, and the single-nuclei ATAC-seq reads were downloaded with ffq and wget from GEO accession GSM6284343 and GSM6284346 respectively.

The ISSAAC-seq human-mouse dataset was obtained from ArrayExpress accession E-MTAB-11264. Reads were downloaded using wget.

### Extracting SHARE-seqV2 barcodes

The SHARE-SeqV2 FASTQ files, for both the RNA and ATAC portion, do not include the nucleotide sequences for the cell barcodes. Barcodes are associated with reads as strings of three numbers placed in the read headers. The numbers point to lines in a file that list the (error-free) 8bp barcodes. Thus, the read headers effectively record the barcodes. Without raw data available, we generated FASTQ files corresponding to the barcode sequences for use with snATAK: the processed barcodes were extracted from each of the FASTQ records and reverse mapped to the barcodes of origin from the known barcode list (https://github.com/pachterlab/BGP_2023/tree/main/references/onlists/shareseq). The three barcodes were then concatenated together with the specific linker structure given by a seqspec specification (Sina Booeshaghi, Chen, and Pachter 2023) to generate a barcode FASTQ file for the ATAC and for the RNA portion of SHARE-SeqV2.

### Comparison of SHARE-seqV2 with ISSAAC-seq

We assessed the multimodal tradeoff between SHARE-seqV2 and ISSAAC-seq separately on human and mouse genes. In order to perform a like-for-like comparison, we subsampled the number of BUS records for the ISSAAC-seq quantifications to match those of the SHARE-seqV2 quantifications. Because the SHARE-seqV2 data was not in raw form with every record containing an already error-free barcode, we subsampled ISSAAC-seq reads after ISSAAC-seq barcode error correction, so that the ISSAAC-seq subsampled data was also no longer raw, and matched the SHARE-seqV2 data exactly in quantity. The average read count per gene was computed for all human genes and all mouse genes. The Pearson correlation was then calculated between ISSAAC-seq and SHARE-seqV2. We also computed the sum of counts for each cell across all human genes and separately for all mouse genes and made a clawplot on the log-log axis (clawplot, Figure 3de,j,k). We summed the number of peaks detected and the number of genes detected for each cell to create the multimodal detection plot. Lastly, using bustools count with the -cm option, we computed the number of read counts per cell per gene.

Then, for each cell, we plotted the sum of the UMI counts vs the sum of the reads and performed linear regression to find a line of best fit. The slope of this line represents the UMI collapse ratio, i.e., how many distinct molecules can be obtained for a given number of reads.

### Downloading Spatial-ATACseq

Reads and count matrices were downloaded from SRA accession GSM5238386 using ffq and wget. The barcode to pixel mapping as well as the list of allowable barcodes was downloaded from GitHub.

### Comparison of Spatial-ATACseq

Reads were processed with snATAK. A pixel-by-pixel matrix of the number of accessible regions per pixel was computed from snATAK quantifications and the spatial-ATAC-seq matrix released on GEO at accession GSE171943. Since the peak count matrices released on GEO contained peaks with overlapping regions we transformed the peaks detected counts to z-scores.

### Downloading 10xPBMC Multiome, snATACseq, and scRNAseq data

10xPBMC Multiome data was downloaded from the 10x Genomics Dataset page for both Chromium X [link] and Chromium Controller [link]. snATACseq data was downloaded from the 10x Genomics Dataset page for both the Chromium X [link] and Chromium Controller [link]. scRNAseq data was downloaded from [link].

### Comparison of Cell Ranger and snATAK on 10x Multiome

We ran Cell Ranger Arc on the 10x Multiome X 10k PBMC dataset and used the peaks and transcriptome reference generated by Cell Ranger with snATAK to assess quantification concordance. ATAC and RNA reads were quantified with snATAK.

### Multiome, snATACseq, and scRNAseq tradeoff

We subsampled all reads to 400 million and then ran standard snATAK to generate count matrices. We then computed the knee plot, peak-peak correlations, and umi-collapse ratio tradeoffs in Figure 2

### Generating Allele-specific count matrices

We downloaded the common_all_20180418.vcf.gz file from the 1k Genomes project and filtered it to obtain a subset of only biallelic SNPs. We further filtered those SNPs to retain SNPs with a minor allele frequency of at least 0.05, and we removed SNPs that were within 30bp of each other. With bedtools, we then created a FASTA file containing a 31 basepair window around each SNP. The FASTA file contained two records per SNP: one for the reference and one for the alternate allele. All records in the FASTA file were also mapped to the genome with BWA fastmap to remove multimapping SNPs. We indexed the remaining SNP FASTA file with kallisto, created a transcripts to gene map of the SNPs and ran snATAK. The gene count matrices were split in Python using scipy and numpy to produce two matrices: one matrix with columns corresponding to the reference allele and one matrix with columns corresponding to the alternate allele.

### Assessing allele concordance

We genotyped the sample by summing the reads across all cells for the alternate and reference SNPs. We performed a simple maximum log-liklihood genotype assignment of homozygous-to-the-reference, heterozygous, or homozygous-to-the-alternate. We selected SNPs that were heterozygous for further analysis.

For each cell type (as found on the RNAseq UMI counts) and for each SNP, we computed the sum of RNA UMI counts for the reference and alternate SNP for the heterozygous SNPs that had either zero ATAC-seq counts in the reference matrix or zero ATACseq counts in the alternate matrix (but not both). To create the SNPs detected plot we computed, for each cell, the number of features with non-zero counts in either ATAC or RNA or both RNA and ATAC.

### Runtime and memory comparison

The /usr/bin/time command was used with the -v option to benchmark runtime and memory for Cell Ranger ATAC, Cell Ranger ARC, and snATAK. Peaks were precomputed and provided to all three programs. Cell Ranger ARC and snATAK were run with 8 threads and 8G max memory options while Cell Ranger ATAC was ran with default parameters.

### Software

The following software tools form the core of the methods constituting the snATAK workflow which was run to generate the results and figures of this paper: kallisto (v0.46.1); bustools (v.0.40.0); minimap2 (v2.15); samtools (v1.10); Genrich; bedtools (v.2.25.0). Notebooks reproducing the results and figures are available at: https://github.com/pachterlab/BGP_2023 In addition to the tools above, the following programs were used for analysis of results and generation of figures:

AnnData 0.7.1 (Virshup et al. 2021) awk (GNU awk) 4.1.4

ffq 0.30 (Gálvez-Merchán et al. 2022) gget 0.27.5 (Luebbert and Pachter 2022) grep (GNU grep) 3.1

kb_python 0.24.4 (Melsted et al. 2021) Matplotlib 3.0.3 (Hunter 2007)

Muon (Bredikhin, Kats, and Stegle 2022) Numpy 1.18.1 (Harris et al. 2020)

Pandas 0.25.3 (Pandas Development Team 2020) sed (GNU sed) 4.4

Scanpy 1.4.5.post3 (Wolf, Angerer, and Theis 2018) Scipy 1.4.1 (Virtanen et al. 2020)

sklearn 0.22.1 (Buitinck et al. 2013)

statsmodels 0.12.1 (Seabold and Perktold 2010) tar (GNU tar) 1.29

### Hardware

All computational work was performed on a Supermicro server computer (2xXeon® Gold 6152 22-Core 2.1, 3.7GHz Turbo, 12 × 64GB Quad-Rank DDR4 2666MHz memory, 16 × 12TB Ultrastar He12 HUH721212ALE600, 7200 RPM, SATA 6Gb/s HDD) with CentOS7 operating system installed.

