## Supplementary Material for "Assessing the multimodal tradeoff"

### Supplementary Information
### Assessing the multimodal tradeoff

A. Sina Booeshaghi[1+], Fan Gao[1,2,+], Lior Pachter[1,3*]

1. Division of Biology and Biological Engineering, California Institute of Technology, Pasadena, CA, USA
2. Caltech Bioinformatics Resource Center, California Institute of Technology, Pasadena, CA, USA
3. Department of Computing & Mathematical Sciences, California Institute of Technology, Pasadena, CA, USA

+These authors contributed equally
*To whom correspondence should be addressed.

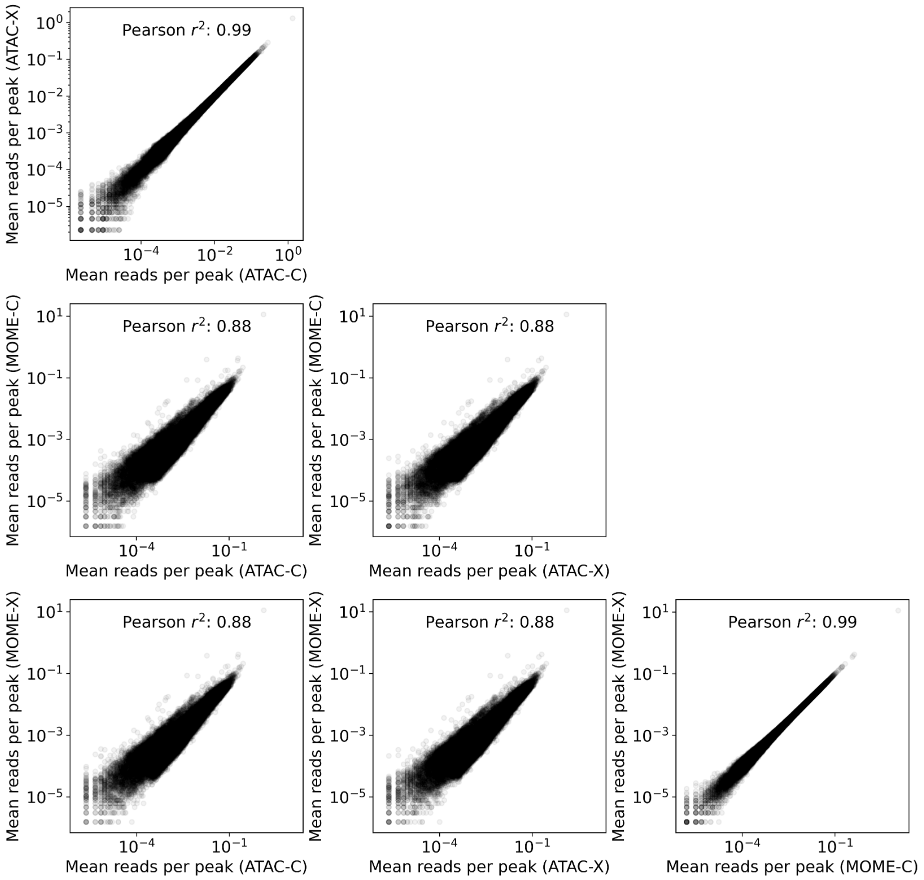


**Supplementary Figure 1:** Peak-peak correlations between four different ATACseq preprocessing assays on 10k PBMCs: 10x Multiome (ATAC) with the Chromium X, 10x Multiome (ATAC) with the Chromium Controller, 10x ATACseq with the Chromium X and 10x ATACseq with the Chromium Controller.

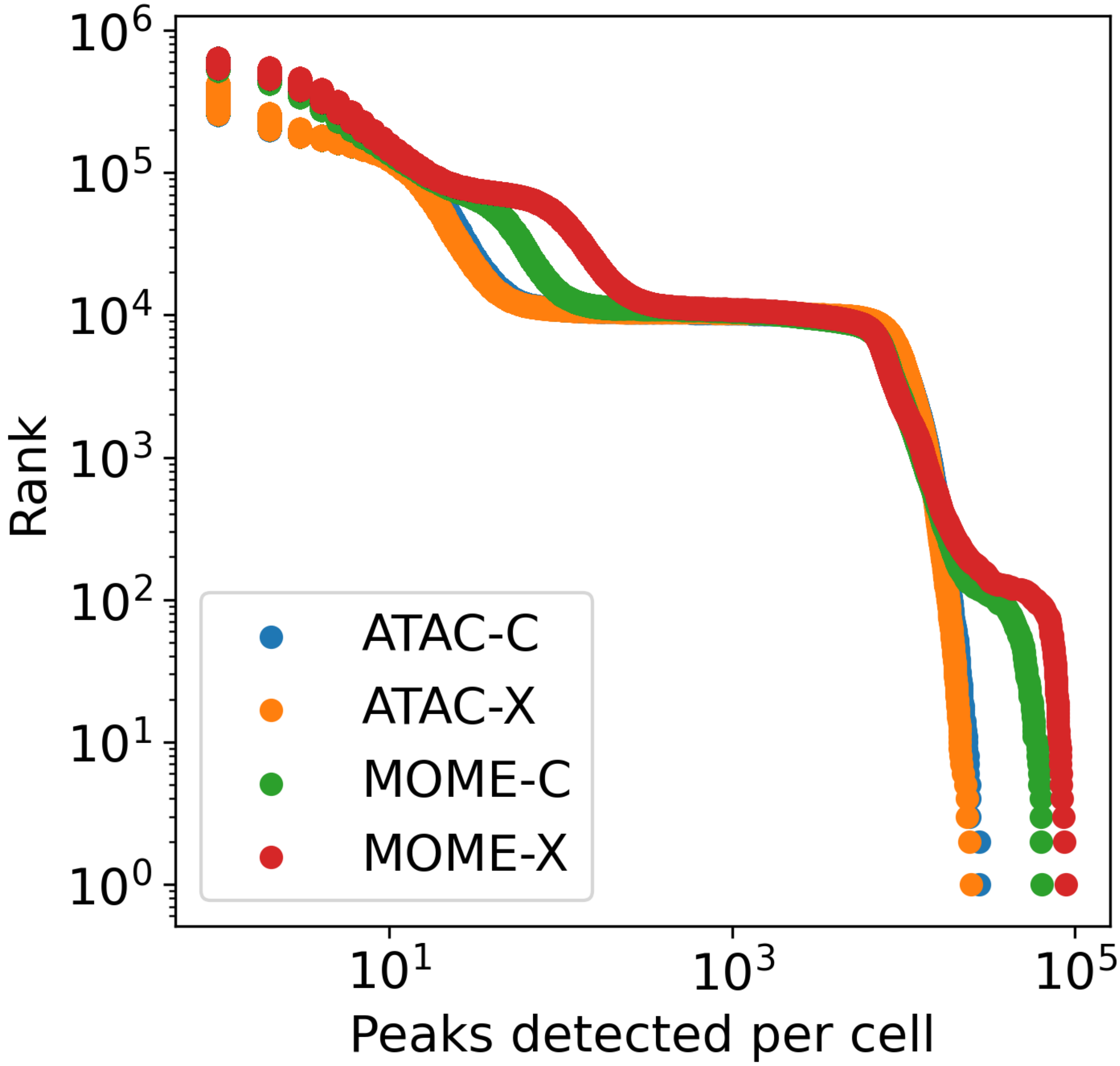


**Supplementary Figure 2:** Reverse cumulative read count distribution per cell for each of the four 10x Genomics ATACseq technologies.

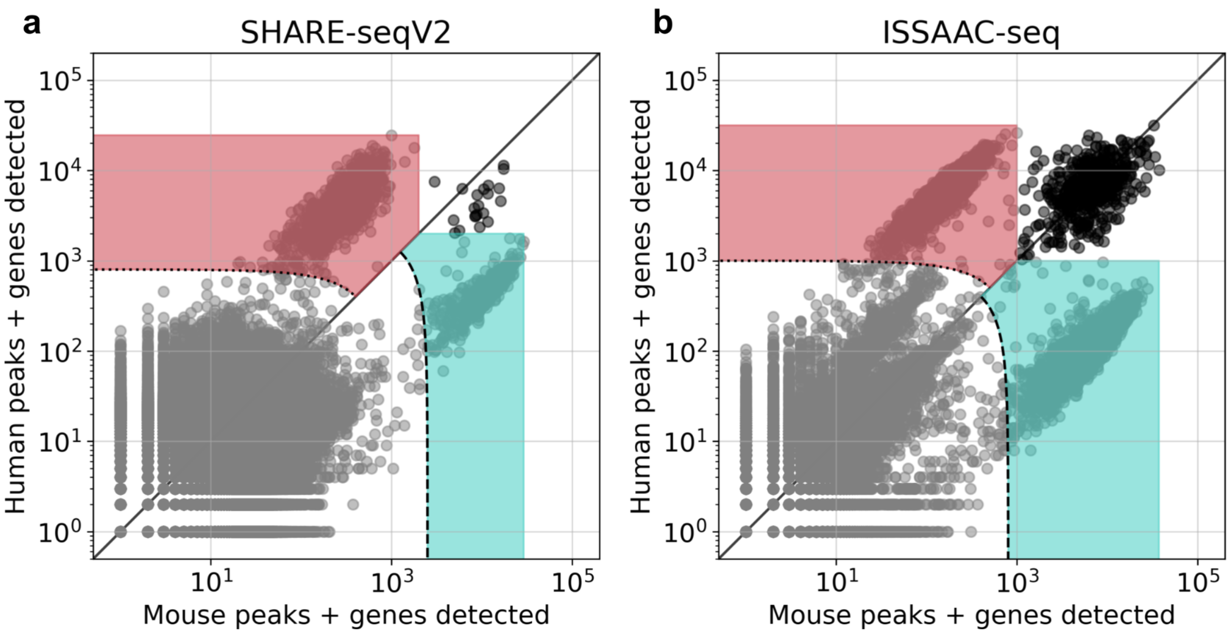


**Supplementary Figure 3:** A comparison of human and mouse genes + peaks detected for SHARE-seqV2 and ISSAAC-seq. The shaded regions in the plots show the selected human and mouse cells, obtained by thresholding low quality cells, and removing multiplets.

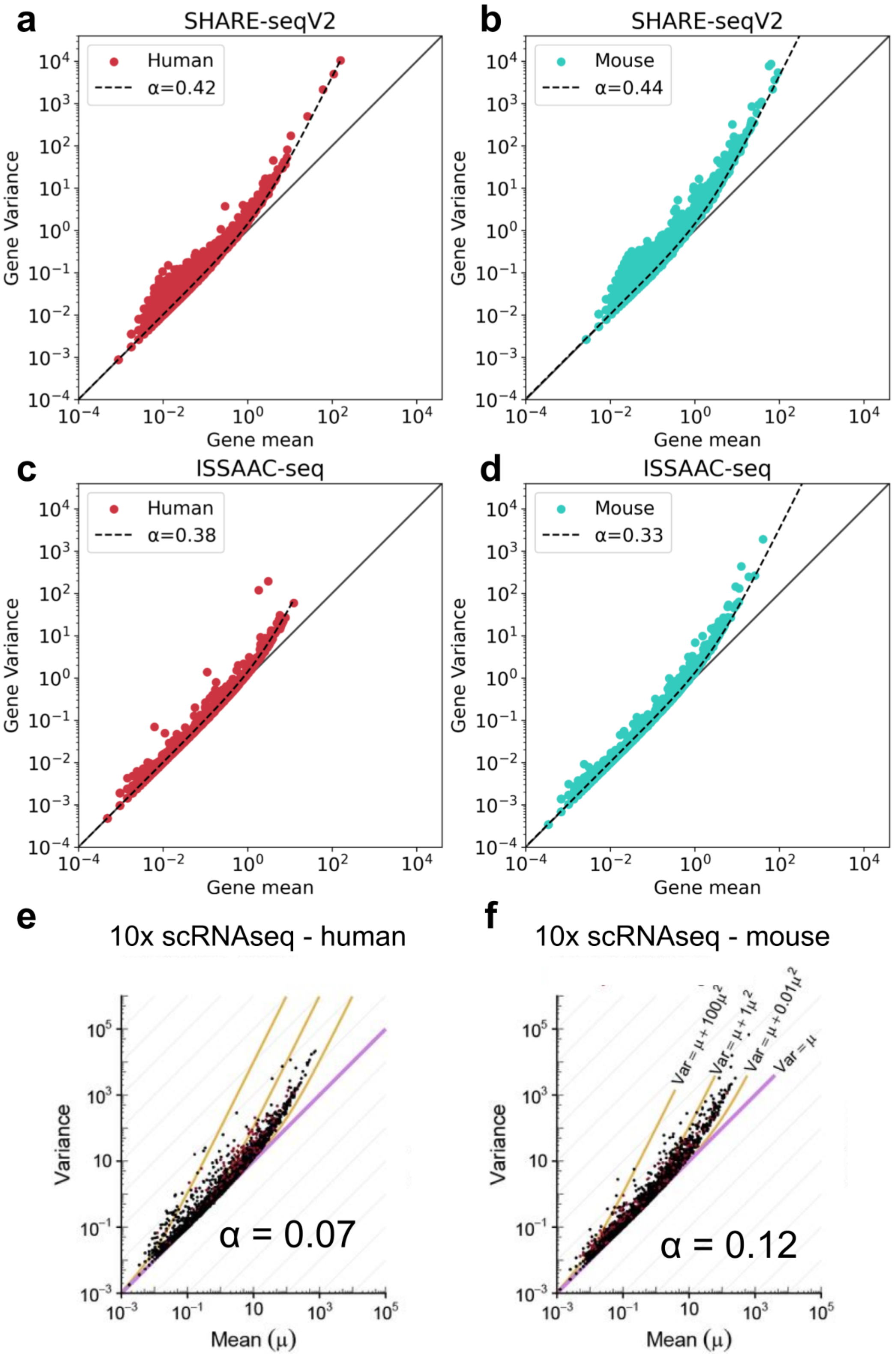


**Supplementary Figure 4:** The mean-variance relationship for human HEK293T cells and mouse NIH3T3 cells: (a) SHARE-seqV2 human, (b) SHARE-SeqV2 mouse, (c) ISSAAC-seq human, (d) ISSAAC-seq mouse, (e) 10x scRNA-seq human (reproduction of part of Figure S1b from (Ahlmann-Eltze and Huber 2021) under the CC-BY-ND 4.0 international license) and f) 10x scRNA-seq mouse (reproduction of part of Figure S1b from (Ahlmann-Eltze and Huber 2021) under the CC-BY-ND 4.0 international license).
